# Climate Adaptation and Genetic Differentiation in the Mosquito Species *Culex tarsalis*

**DOI:** 10.1101/2024.09.23.613315

**Authors:** Yunfei Liao, Touhid Islam, Rooksana Noorai, Jared Streich, Christopher Saski, Lee W. Cohnstaedt, Elizabeth A. Cooper

## Abstract

The increasing prevalence of vector-borne diseases around the world highlights the pressing need for an in-depth exploration of the genetic and environmental factors that shape the adaptability and widespread distribution of mosquito populations. This research focuses on *Culex tarsalis*, a principal vector for various viral diseases including West Nile Virus (WNV). Through the development of a new reference genome and the examination of Restriction-Site Associated DNA sequencing (RAD-seq) data from over 300 individuals and 28 locations, we demonstrate that variables such as temperature, evaporation rates, and the density of vegetation significantly impact the genetic makeup of *Cx. tarsalis* populations. Among the alleles most strongly associated with environmental factors is a nonsynonymous mutation in a key gene related to circadian rhythms. These results offer new insights into the mechanisms of spread and adaptation in a key North American vector species, which is poised to become a growing health threat to both humans and animals in the face of ongoing climate change.

## Introduction

Diseases transmitted by insects are a serious and growing health concern for both humans and livestock. Numerous mosquito species are viral vectors, and the global distribution plus high adaptability of these mosquitoes contributes to the rapid spread and evolution of dangerous diseases across multiple continents (Gubler 1998). Understanding how different species of mosquito have increased their ranges and adapted to new habitats in the past will be critical for predicting and managing their potential expansion in the future (Githeko et al. 2000; Sutherst 2004).

The most abundant vector species found in the United States are derived from two genera: *Culex* and *Aedes*, each of which is comprised of both native and introduced species that occupy a variety of geographical ranges (Darsie and Ward 2016). *Culex tarsalis*, also known as the Western Encephalitis Mosquito, is a carrier of several forms of encephalitis that can infect humans as well as animals such as horses(Reeves 1990; Darsie and Ward 2016). It is a known vector of West Nile Virus (WNV), Japanese Encephalitis Virus, St. Louis Encephalitis Virus, and Rift Valley Fever Virus (Evans et al. 2017; Main et al. 2018). *Cx. tarsalis* is abundant in the western continental United States, where it is responsible for the majority of WNV cases in the most severely impacted states (Goddard et al. 2002; Evans et al. 2017).

Despite the economic and health risks posed by this mosquito species, there is little known about its genetics. Previous studies using microsatellite markers have consistently uncovered a distinct pattern of population structure that does not entirely correlate with current geographical features or indicate strong isolation-by-distance (Venkatesan and Rasgon 2010; Pfeiler et al. 2013), but genetic results do support the hypothesis that *Cx. tarsalis* originated on the southwest coast of North America and has since undergone a range expansion to spread eastward (Venkatesan et al. 2007). While significant adaptive changes must have occurred to allow populations to overwinter in order to cross the Rocky Mountains (Venkatesan et al. 2007; Diniz et al. 2017), the precise geographic and climatic variables driving divergence among populations are largely unknown.

Although the pattern of genetic differentiation observed in *Cx. tarsalis* could be the result of historical geographic divisions, it is also possible that local environmental adaptations may be driving some or all the population divergence in this species. Many mosquito species, including other *Culex* species, exhibit little to no population structure even after short time spans due to their large population sizes and short generation times (Wilke et al. 2014; Kotsakiozi et al. 2017), so the pattern observed in *Cx. tarsalis* is atypical and suggests a potential role for selection in addition to genetic drift. If the range of present-day populations of *Cx. tarsalis* is defined by adaptation to certain environmental factors, then identifying these factors as well as the genes and alleles under selection is essential to predicting whether or not *Cx. tarsalis* could continue to spread eastward and northward while being a more prevalent threat within the United States and possibly even other countries.

To advance our understanding of population structure and identify alleles linked to local adaptation in *Cx. tarsalis*, we first assembled and annotated a *de novo* reference genome and generated Restriction-Site Associated DNA sequencing (RAD-seq) data for over 300 individuals from 28 diverse geographic locations. We analyzed these RAD-seq markers through a comprehensive landscape genetics framework to explore how various environmental variables influence population differentiation and to identify alleles associated with adaptation to these conditions. By leveraging a broad spectrum of environmental variables, we assessed the adaptive responses of populations to their local environments, enabling the identification of critical genetic-environment associations. This approach reveals how specific climate variables and genetic variants underpin local adaptation strategies across 28 representative *Cx. tarsalis* collection sites. Our findings enrich our understanding of the complex interactions between genetics and environment, providing crucial insights into the ecological dynamics of this mosquito species.

## Results

### Genomic Analyses

The final genome assembly contained 968,887,694 bases divided into 7,478 contigs. The N50 was 451,230 bp. The annotation included 43,905 predicted genes. After aligning the RAD-seq reads and filtering for quality, there were 457,387 polymorphic sites identified across all populations.

### Population Structure

The ADMIXTURE analysis indicated a strong signature of population structure among the collected samples, with the optimal number of population assignments occurring at K=4 (Supplemental Figure 1). The genetic clusters corresponded to four different broad geographic regions: (1) California/the West Coast, (2) the Southwest, (3) the Northwest, and (4) the Midwest (Figure 1).

**Figure 1:**
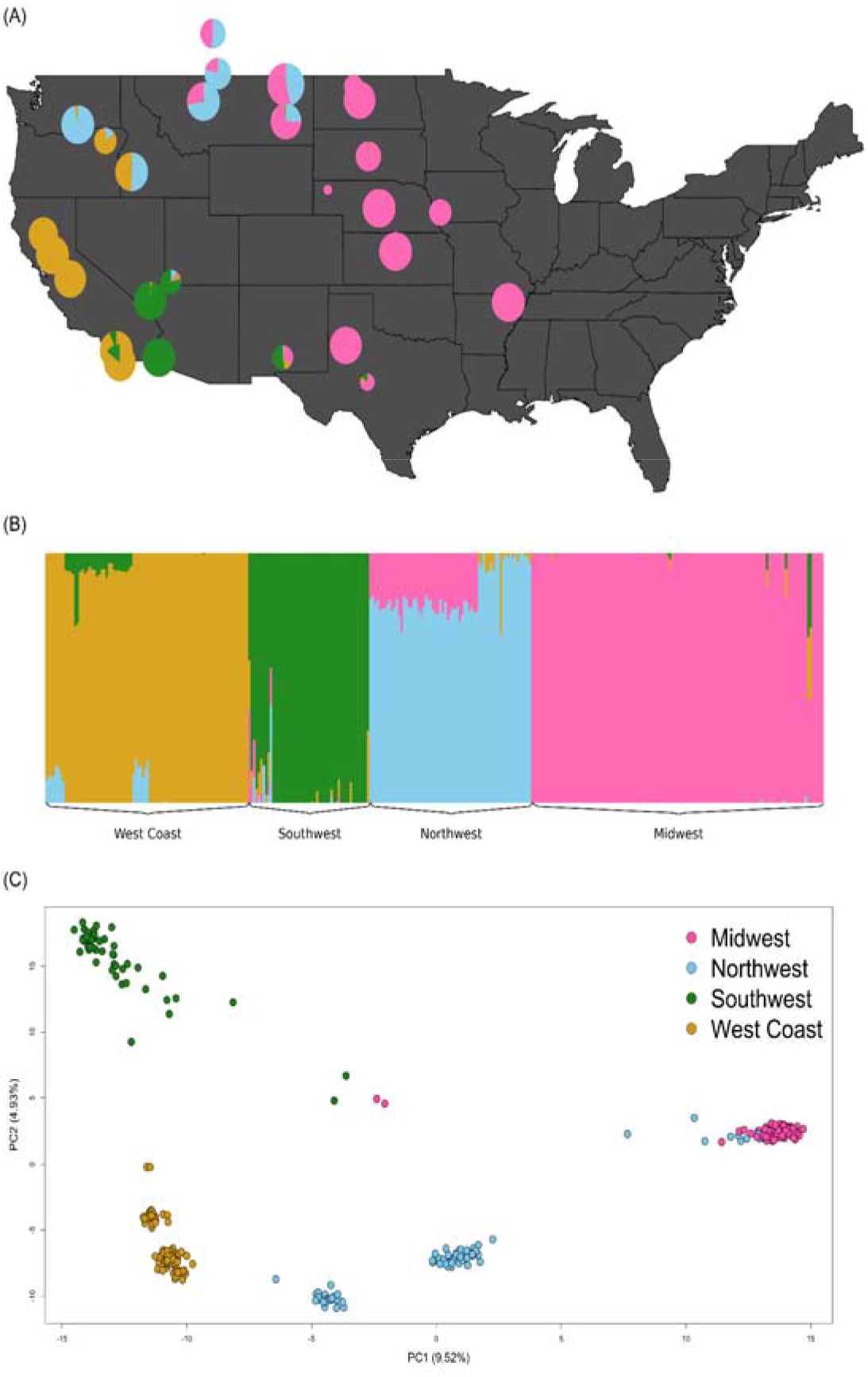
Population Structure of *Cx. tarsalis*: (A) Floating pie charts of the admixture proportions in *Cx. tarsalis* populations sampled across the Western and Midwestern U.S and parts of Canada. Pie chart sizes are proportional to the sample size at each collection site. (B) ADMIXTURE results for K=4. Labels along the x-axis indicate sampling locations and colors correspond to the admixture proportion for each of the 4 clusters. (C) PCA results for the top 2 principal components, with points colored by the 4 geographic regions identified by ADMIXTURE.

The PCA results confirmed this pattern (Supplemental Figure 1 and Figure 1), while also showing evidence of some sub-structure among the West Coast and Northwest populations (Figure 1C).

The AMOVA results also indicated significant levels of population differentiation both between different populations within the same geographic region, and between different regions (Supplemental Table 1 and Figure 2). The observed genetic variation within populations was significantly lower (p < 0.001) than expected (Supplemental Figure 2a and Supplemental Table 2), while variation between populations and between regions was significantly higher (p < 0.001) than would be expected by chance (Supplemental Figure 2b and 2c, Supplemental Table 2).

**Figure 2:**
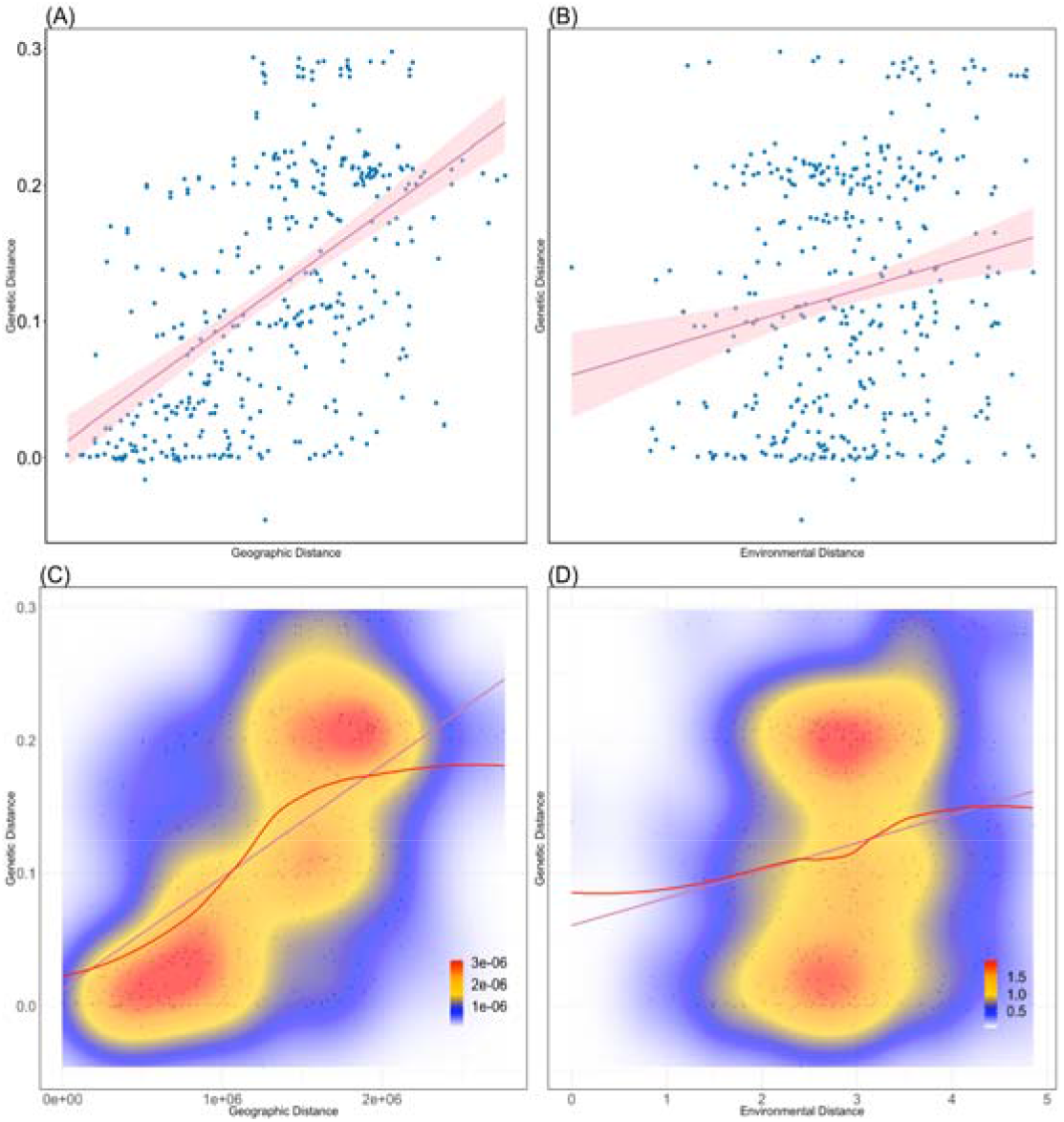
Isolation-by-Distance and Isolation-by-Environment. (A) Pairwise geographic distance versus genetic distance (F_ST_) with best fit linear regression model (red line:). (B) Pairwise environmental distance versus genetic distance with best-fit linear regression mode (red line:). (C)Kernel density plot with best fit spline for geographic distance versus genetic distance. Areas of high, intermediate, and low density are represented by red, yellow, and blue colors, respectively. (D) Kernel density plot with best-fit spline for environmental distance versus genetic distance.

### Isolation by Distance (IBD) verse Isolation by Environment (IBE)

Both geographic distance and environmental distance showed a statistically significant relationship with genetic distance (Figure 2). The correlation with geographic distance (IBD) was much stronger, with a Mantel test statistic of 0.5661 (p=0.001), while the correlation with environmental distance (IBE) was weak but still significant, with a Mantel test statistic of 0.1866 (p=0.0115). The mixed model results indicated that considering both geographic and environmental factors simultaneously provided a more comprehensive explanation of genetic distance variations than either factor alone (Table 1). The moderate to strong correlation observed in IBD and the weaker correlation in IBE are echoed in the mixed model, where the combined effect of both distances was most significant in explaining genetic differences. These results suggest that while both geographic and environmental factors contribute to genetic differentiation, geographic factors play a more dominant role in the system under study. Further, the kernel density plot and LOESS fit in Figure 2C suggest a non-linear relationship between geographic and genetic distances, implying variability in correlation across different ranges and the potential influence of complex factors rather than simple geographic separation. Similarly, in Figure 2D, a weaker and more uniform correlation between genetic and environmental distances is shown, with a lower R² of 0.04, where the LOESS fit line closely follows the linear regression with minor deviations at smaller ranges. This pattern supports the idea that other influencing elements may affect the relationship, which may not be solely decided by linear factors.

**Table 1:** Mixed model results for IBD and IBE.

| Models | AIC | BIC | k | AICc | AICcmin | BICew |
| --- | --- | --- | --- | --- | --- | --- |
| Full | 689.701 | 709.375 | 5 | 689.862 | 0.959 | 0.771 |
| Distance | 696.061 | 711.801 | 4 | 696.169 | 0.041 | 0.229 |
| Environment | 1001.974 | 1017.713 | 4 | 1002.081 | 0.000 | 0.000 |

### Genotype-Environment Associations

In the PCA of the 8 environmental variables, described in the Materials and Methods section, the first component explained 34.75% of the variation in the predictors, with the second component accounting for 21.42% of the variance, and the third component accounting for 16.73%, for a total cumulative explained variance of 72.9% (Supplemental Table 3 and Supplemental Table 4). Using the first PC as the predictor in the LFMM, 92 candidate SNPs were identified as being significantly associated with the environment (FDR < 0.1) (Supplemental Table 5).

The RDA model revealed that environmental variables explain 10.42% of the genetic variance (Constrained), leaving 89.58% of the variance unexplained and presumably accounted for by geographical distance as supported by the IBD and IBE tests, the environmental contribution to genetic variance nonetheless provides valuable insights into adaptive processes. Even though the majority of the variance was not explained by environmental factors, both the overall RDA model and the first 4 axes still demonstrated statistical significance, indicating that there were substantive associations within the data, which suggests the presence of meaningful genotype-environment associations (Table 2). A total of 822 SNP candidates were detected as significantly associated with at least one of the first four RDA loadings (Supplemental Table 6). Among these, 658 were found within genes. Additionally, out of all the genes identified by RDA containing the 658 SNPs, 32 genes were also found to be significantly associated with the environment in the LFMM (Figure 5).

**Table 2:** Significant test results on each constrained axis of RDA.

|  | Df | Variance | F | Pr(>F) | Significance |
| --- | --- | --- | --- | --- | --- |
| Full Model | 8 | 1796.3 | 4.5511 | 0.001 | *** |
| RDA1 | 1 | 894.1 | 18.1217 | 0.001 | *** |
| RDA2 | 1 | 469.8 | 9.5231 | 0.001 | *** |
| RDA3 | 1 | 158.7 | 3.2158 | 0.001 | *** |
| RDA4 | 1 | 69.8 | 1.4154 | 0.001 | *** |
| RDA5 | 1 | 56.8 | 1.1517 | 0.094 | . |
| RDA6 | 1 | 51.1 | 1.0361 | 0.591 |  |
| Full Model | 8 | 1796.3 | 4.5511 | 0.001 | *** |
| RDA1 | 1 | 894.1 | 18.1217 | 0.001 | *** |
| RDA2 | 1 | 469.8 | 9.5231 | 0.001 | *** |
| RDA7 | 1 | 48.6 | 0.9855 | 0.901 |  |
| RDA8 | 1 | 47.3 | 0.9592 | 0.901 |  |
| Constrained axis Residual | 313 | 15442.7 |  |  |  |

Significant Code: 0 '\*\*\*' 0.001 '\*\*' 0.01: '\*' 0.05 '.' 0.1 ' ' 1

Figure 3A captures the distribution of mosquito populations across the RDA 1 and RDA 2 axes, which together explained 56% of the environmentally influenced genetic variance (corresponding to 5.84% of the total genetic variance). The clear regional clustering depicted within this biplot aligns with the four geographic regions (Midwest, Northwest, Southwest, and West Coast), underscoring a significant regional influence on the portion of genetic variation shaped by environmental factors. Specifically, the alignment of populations with vectors for the temperature and evaporation signifies the role of temperature and humidity in this context. Figure 3B further explores the subtler environmental gradients within the RDA 3 and RDA 4 axes, which together explain approximately 15% of the environmentally responsive genetic variance (equating to roughly 1.56% of the total genetic variance). Here, the distributions suggest a more intricate interaction of genetic variance with environmental variables like the surface runoff and leaf area index for high vegetation, which could reflect micro-environmental adaptations.

**Figure 3.**
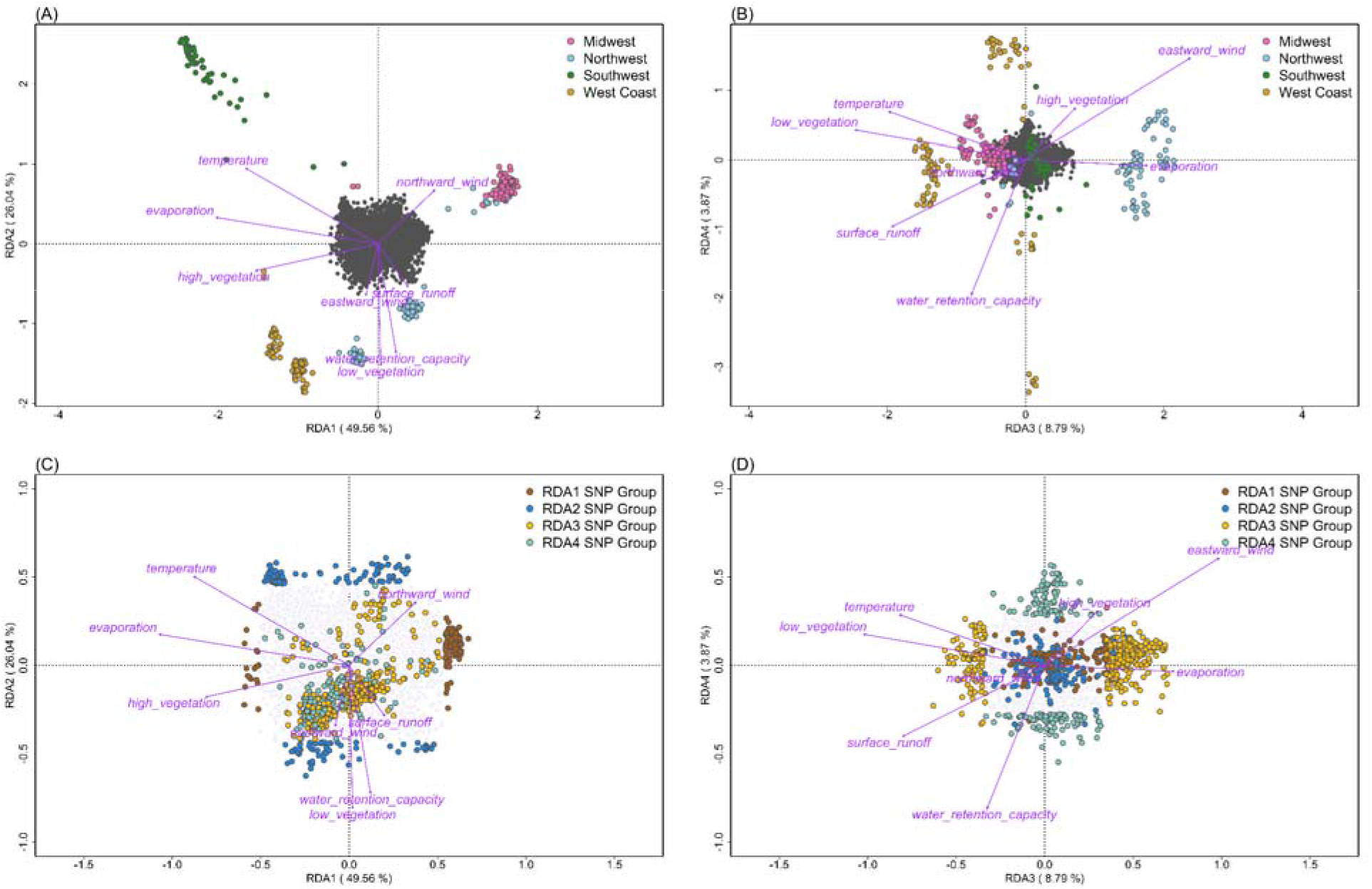
Environmental Correlates of Genetic Variation in *Cx. tarsalis* Across Diverse North American Regions. Panels A and B display the relationship between environmental factors and the distribution of *Cx. tarsalis*, using RDA to illustrate how regional differences affect genetic variation. In these panels, the position of each circle (representing an individual mosquito) and color (indicating regional groupings from ADMIXTURE results) reflects their association with environmental variables, shown as purple vectors. The first plot (A) focuses on RDA1 and RDA2, the primary axes explaining the most variance, while the second (B) explores more subtle influences in RDA3 and RDA4. Panels C and D shift focus to SNPs within the genetic data, colored to show which RDA constrained axis they are extracted. These latter plots further detail the genetic-environment relationship, with the SNPs’ distribution providing insight into the adaptive landscape of these populations.

In Figure 3C, the focus shifts to candidate SNPs, arrayed along the RDA 1 and RDA 2 axes and color-coded by which RDA constrained axis they are extracted from. The majority of the candidate SNPs in the RDA1 group are negatively correlated to evaporation, temperature and high vegetation, aligning with Figure 4 and Supplemental Table 7. As shown in Figure 3A, the first and fourth quadrants likely represent eastern populations, while the second and third quadrants represent western populations. This east-west gradient is evident among the RDA1 SNPs group. Similarly, the RDA2 group candidate SNPs are more negatively correlated to low vegetation index, water retention capacity but positively correlated to temperature. Figure 3D’s examination of the RDA 3 and RDA 4 axes along with the result in Supplemental Table 7 provides a layered view of SNP distribution. This analysis reveals a different pattern, with eastward wind and surface runoff showing stronger correlations than in the RDA1 and RDA2 groups. As a result, most significantly associated SNPs were correlated with more than one environmental variable, and often clustered together in distinct patterns (Figures 4c and 4d, Figure 4). Although this complex interaction represents a smaller slice of the total genetic variance, it is critical for a comprehensive understanding of genetic-environment relationships.

**Figure 4.**
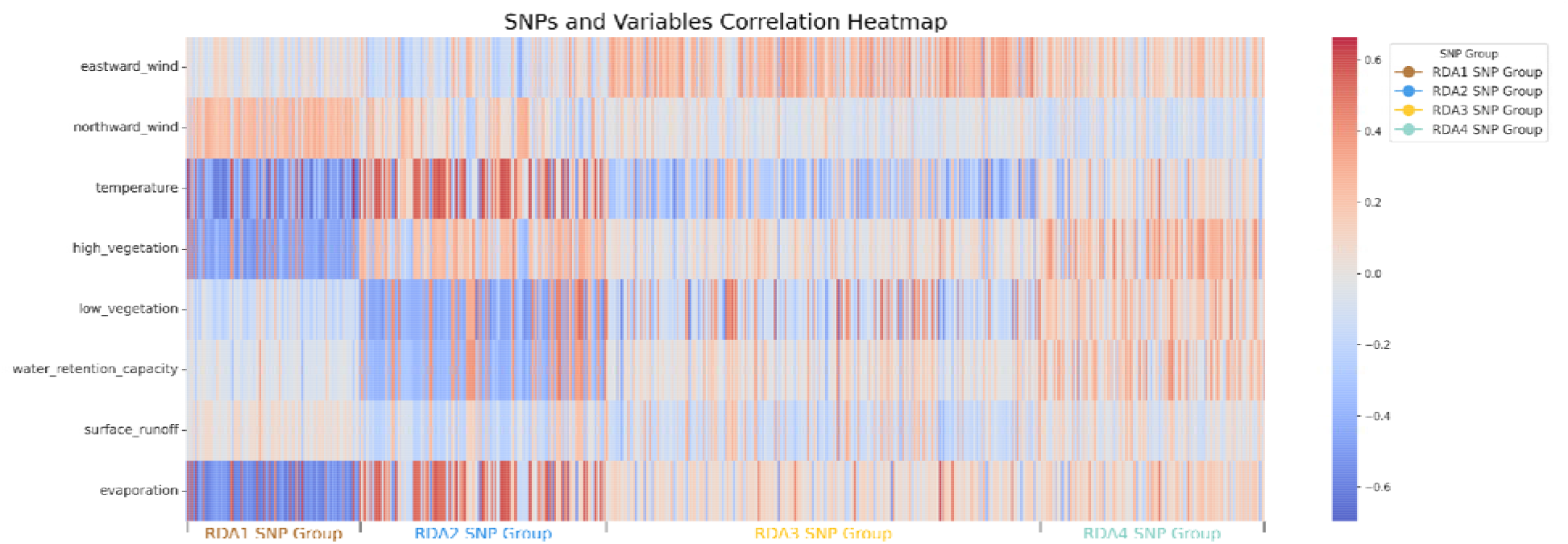
Correlation Heatmap between Environmental Variables and RDA Candidate SNPs. This heatmap illustrates the correlation between the environmental variables and candidate SNPs identified through the first four constrained axes of Redundancy Analysis (RDA). Each column on the x-axis represents a candidate SNP. For detailed SNP names and associated information on the 17 genes identified by SIFT4G, please refer to Supplemental Figure 3.

This suite of RDA plots collectively reveals the environmental portion of genetic variance within *Cx. tarsalis* populations. While most of the genetic variation correlates with geographic distribution, the environmental variance captured here affords a critical perspective on the selective forces at play, contributing to the broader evolutionary narrative of this species.

Overall, our findings underscore the intricate connections between geographic locations, environmental factors, and genetic variations, confirming that in addition to genetic drift and IBD, there is a profound influence of environmental variables on genetic variation in *Cx. tarsalis*.

### SNPs Under Selection

The BayeScan outlier analysis revealed 1,836 loci potentially under diversifying selection, 10,166 potentially under balancing selection, and 5,237 neutral loci (Supplemental Figure 4 and Supplemental Table 8). Of the 1,836 loci putatively undergoing selection for local adaptation, 1501 of these are found within 824 genes. Of these 824 genes, 24 also overlapped with genes that were identified by both the LFMM and RDA as significantly associated with environmental variables. We also independently used PCAdapt to search for outlier loci with notable allele frequency differences across populations that are potentially the result of natural selection and uncovered 173 SNPs in the top 1% of extreme p-values (Supplemental Figure 5 and Supplemental Table 9). In total, 53 candidate local adaptation SNPs were identified (Supplemental Table 10). Some candidates show a strong East-West gradient in terms of allele frequencies (Supplemental Figure 6A and 6B), while others indicate that the alternate allele is present only in one or a few populations (Supplemental Figure 6C and 6D). Looking across all 4 of our analyses examining significant environmental associations and evidence of natural selection, we found 20 common genes that contain the candidate SNPs that were significant in every instance (Figure 5 and Supplemental Table 11). A Chi-Square test (Supplemental Table 12 and Supplemental Table 13) indicated a significant association between the candidacy of SNPs and their location being within genes rather than intergenic (X-squared = 75.842, df = 1, p-value < 2.2e-16). This suggests that SNPs identified as candidates are more likely to be located within genes than would be expected by chance.

**Figure 5:**
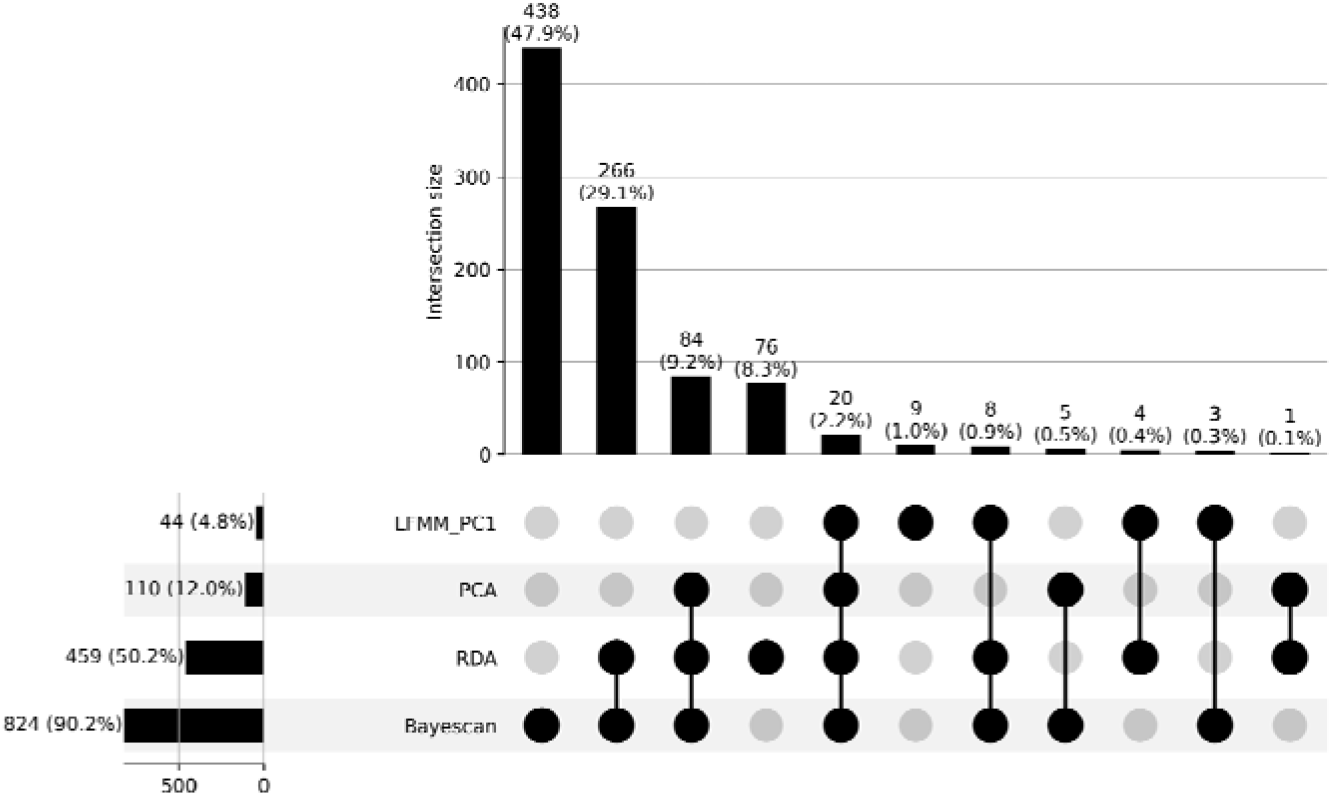
Upset Plot of within Genes Annotated from Candidate SNPs Identified for Local Adaptation and Environmental Association Across LFMM, RDA, Pcadapt, and BayeScan Analysis in *Cx. tarsalis*.

The SIFT4G predictions determined that 17 of the 20 overlapping candidate genes contained an environmentally-associated SNP located within the coding region, and 8 of these genes contained nonsynonymous mutations within our dataset (Supplemental Table 10). Among these genes, several genes are noteworthy due to their intriguing functions. For instance, Ct.00g030230, identified as a seminal plasma protein, may play a role in reproductive success (Boes et al. 2014; Amaro et al. 2021). Ct.00g095350, a defective proboscis extension response-related (DPR) protein has been shown to affect feeding behavior in fruit flies (Nakamura et al. 2002). Ct.00g154760, which was annotated as a carnitine O-acyltransferase, is involved in fatty acid metabolism (Jogl et al. 2004), a pathway crucial for energy provision under varying climatic conditions. Lastly, Ct.00g04290 encodes a PERIOD CIRCADIAN PROTEIN, which is essential for the regulation of circadian rhythms, aligning life cycle events with environmental cues (Meuti et al. 2015; Chang and Meuti 2020; Shetty et al. 2022). Notably, within this gene, two nonsynonymous and one synonymous SNPs were discerned. SIFT predictions indicated that one of the nonsynonymous SNPs could be deleterious, potentially impacting the protein’s functionality and, by extension, the organism’s adaptability to environmental rhythmic changes. This was the one of the two candidate SNPs classified as deleterious by SIFT, so it represents a significant point of interest for its potential role in the ecological adaptation of *Cx. tarsalis*.

## Discussion

We successfully assembled a draft genome for *Cx. tarsalis* and identified hundreds of thousands of polymorphic markers across individuals sampled from diverse locations in North America, primarily in four regions: the West Coast, Southwest, Northwest, and Midwest. Each region is characterized by unique climatic conditions that likely influence genetic variation. The West Coast, known for its warm, seasonally dry climate, contrasts quite significantly with the hot, dry conditions and sparse vegetation of the Southwest. In comparison, the Northwest features a cool, wet climate with lush high vegetation, while the Midwest has a cool, dry climate characterized by a mix of both high and low vegetation. These data revealed a distinct pattern of population structure within this species, with clear differentiation among populations from these regions. Additionally, there was a strong correlation between geographic and genetic distances, highlighting the roles of genetic drift and selection in shaping genetic variation across these varied landscapes.

Our analysis also uncovered a significant link between environmental variables and genetic variation, particularly showing that temperature, evaporation rates, and vegetation density are critical environmental factors with strong associations to genetic differentiation within *Cx. tarsalis* populations. Among these, the identification of 53 SNPs with strong evidence of both selection and environmental correlation across multiple tests suggests that these are most likely to be involved in local adaptation processes. These SNPs, linked to crucial biological functions such as circadian rhythms, reproductive success, feeding habits and fat metabolism, and, lay the groundwork for a detailed exploration of the genetic mechanisms driving adaptation in diverse environmental conditions.

Probably the most interesting SNP uncovered in our study was a deleterious mutation in a gene that we found to be the single copy ortholog of per, a well-studied circadian rhythm gene that encodes the regulatory period protein(Konopka and Benzer 1971). Circadian clock genes are critical for synchronizing physiological and behavioral processes in insects as well as nearly all other living organisms, so a mutation in this gene could be key to the mechanisms that have allowed *Cx. tarsalis* to expand both northward and eastward across the North American continent. Research in other vector species have found that variations in circadian clock genes have a profound impact on mosquito behavior and fitness in both *Cx. pipiens* (Meuti et al. 2015; Chang and Meuti 2020) and the more distantly related Aedes aegypti (Shetty et al. 2022), and in *Cx. pipiens* it was found that circadian regulators (including per) were necessary for inducing diapause.

While it has been observed that at least some *Cx. tarsalis* populations must be capable of entering diapause in order to navigate the challenges of seasonal extremes and to traverse significant geographical barriers, such as the Rocky Mountains, the genetic mechanisms underlying this trait have not yet been identified in *Cx. tarsalis*. Our discovery of a candidate allele in the per ortholog represents the first identification of a potential genetic mechanism governing diapause in this species. The elucidation of circadian protein-involved feedback loops, as explored by Shetty et al., further emphasizes the potential universality of these mechanisms across mosquito species. This breakthrough underscores this gene’s central role in broader adaptive strategies crucial for ecological adaptation in insect populations.

In addition to the circadian regulator gene, we also identified a handful of other highly significant nonsynonymous mutations in other genes related to traits that are known to be key for insect survival during seasonal changes. For example, Ct.00g030230 was identified as encoding a seminal plasma protein, a family of proteins which are known to influence fertility and post-mating behavior in mosquitoes (Boes et al. 2014; Amaro et al. 2021). Ct.00g095350 was homologous to the DPR gene in Drosophila, where it has been shown to have a role in regulating feeding behavior (Nakamura et al. 2002).

In fruit flies, the DPR gene is involved in the gustatory (taste) response, particularly in the aversion to salt. In mosquitoes, feeding behavior is critical for both nutrient intake and disease transmission, so this gene may be crucial for not only understanding the ecology of *Cx. tarsalis*, but also its disease transmission dynamics. Finally, we also found a significant mutation in the gene Ct.00g154760, which encodes a carnitine O-acyltransferase. Carnitine O-acyltransferase facilitates the transport of fatty acids into mitochondria for β-oxidation, a process integral to energy production (Jogl et al. 2004). Adaptations in Ct.00g154760 that enhance the enzyme’s efficiency could provide *Cx. tarsalis* with survival advantages by optimizing energy utilization under fluctuating conditions, and could also be related to diapause behaviors, since insects must accumulate fat reserves prior to entering into the diapause state(Denlinger 2002).

Overall, our investigation into the *Cx. tarsalis* genome has highlighted the critical importance of environmental adaptations for understanding this mosquito’s distribution and adaptation to a wide range of habitats across North America. Identifying genetic markers linked to circadian rhythms, reproductive processes, and metabolic functions reveals how selection on a few key genetic variants may have sculpted the species’ ability to adapt to an array of environmental challenges and expand its range both northward and eastward. Such genetic insights are pivotal as climate change continues to reshape habitats, potentially enabling vector species like *Cx. tarsalis* to spread into new areas and present new health challenges in the near future.

## Materials and Methods

### Sample Collection

Individual mosquitoes were trapped and collected from 28 different locations across the United States and Canada as part of the North American Mosquito Project (NAMP) (Cohnstaedt et al. 2016). All samples used in this study were collected in 2012 between the months of April and October.

### Genome Sequencing, Assembly, and Annotation

An F4 population was used to generate the reference genome assembly, and high molecular weight DNA was extracted and sequenced on a Pacific Biosciences (PacBio) RS II (University of Delaware). Thirty-five SMRTcells were generated. The resulting reads provided 76X coverage of the ∼790Mb *Cx. tarsalis* genome, and were assembled with MECAT (Xiao et al. 2017).

Gene annotation was completed by MAKER (Cantarel et al. 2008) using EST and protein data from the *Culex quinquefasciatus* and *Aedes aegypti* mosquitoes. Sequences were downloaded from the NCBI Taxonomy database and both Trinotate and InterProScan were used for functional annotation of the MAKER predicted genes (Jones et al. 2014; Bryant et al. 2017). The annotated assembly was assessed for completeness and quality using BUSCO (Seppey et al. 2019) and QUAST (Gurevich et al. 2013).

### RAD-Seq Library Preparation, Sequencing, and SNP Calling

DNA was extracted from individual mosquitoes and libraries were constructed for Restriction-site Associated DNA Sequencing (RAD-Seq) according to previously established protocols (Etter et al. 2011). The SbfI enzyme was used to digest purified DNA, and individual samples were barcoded prior to Illumina sequencing. Raw sequencing reads were subsequently filtered to remove any reads with an uncalled base, an error in the restriction enzyme cut site, or with an average Phred quality score less than 20 over 15 consecutive nucleotides. Filtered reads were then de-multiplexed using the Stacks software package (Etter et al. 2011; Catchen et al. 2013).

After de-multiplexing, raw reads from each individual were aligned to the draft assembly of the *Cx. tarsalis* genome using BWA MEM (Li and Durbin 2009), and individuals with poor mapping rates (less than 50%) were excluded from subsequent analyses. The mapped reads for the remaining 378 samples were then merged using the Samtools pipeline (H. Li et al. 2009) and SNPs were called using the GATK HaplotypeCaller (McKenna et al. 2010). The SNPs were filtered using VCFtools v0.1.12a (McKenna et al. 2010; Danecek et al. 2011) to retain only sites with a minimum average individual read depth of 10X and a maximum of 20% missing data, resulting in a total of 457,387 sites. Individual samples were then filtered again to remove individuals with missing data at more than 50% of the remaining SNP sites, leaving 322 samples from 28 different locations for further analysis. Coding and noncoding SNP effects were predicted using SIFT4G (Vaser et al. 2016).

### Population Structure Analyses

For population structure analyses, SNPs were further filtered using VCFtools v0.1.16 (Danecek et al. 2011) to retain only bi-allelic sites with a maximum of 20% missing data per site and a minimum minor allele frequency (MAF) of 5%. The ADMIXTURE v1.3.0 program (Alexander et al. 2009) was subsequently employed on the remaining 17,239 loci and 322 individuals. Multiple runs of ADMIXTURE were conducted for K values ranging from 1 to 13, with each K value analyzed across 10 independent runs using different random number seeds in order to ensure convergence. The best K value was determined through a cross-validation process ranging from K=2 to K=13 (Supplemental Figure 1). Subsequently, ancestry proportion bar plots were generated using Python’s pandas library (plot.bar()), and floating pie charts displaying ancestry proportions over the U.S. map were plotted using the ggplot2 package in R (Villanueva and Chen 2019) along with scatterpie() in the scatterpie package (Yu).

Complementing this, a Principal Component Analysis (PCA) was performed in R to aid in determining the pattern of genetic differentiation and the optimal K value by visualizing the dataset in a reduced dimensional space. As PCA requires no missing data, the missing genotype values for a given SNP were imputed using the most common genotype at each SNP across all individuals (Y. Li et al. 2009).

An Analysis of MOlecular VAriance (AMOVA) (Excoffier et al. 1992) was performed using the poppr package’s poppr.amova() function in R (Kamvar et al. 2014) (Excoffier et al. 1992) to detect population differentiation among regions inferred from ADMIXTURE results. Pairwise FST values between all populations were computed using the hierfstat package’s genet.dist() function in R (Goudet 2005), employing the Weir-Cockerham estimator (Weir and Cockerham 1984). To assess whether there was a statistically significant level of population structure, a randomization test was conducted using the randtest() function from the ade4 package in R (Thioulouse et al. 2018), employing 999 replicates.

To analyze isolation by distance (IBD) (Slatkin 1993) and isolation by environment (IBE) (Wang and Bradburd 2014; Jiang et al. 2019) patterns within the 28 mosquito populations, genetic distances, derived from Weir & Cockerham FST estimations (Weir and Cockerham 1984) based on SNP data, were compared with geographic and environmental distances computed based on latitude and longitude coordinates. Environmental distances were calculated in R using the dist() function, applying the Canberra method to emphasize relative differences in 8 non-negative environmental variables sourced from the Copernicus Climate Change Service for each geographic location. Scatterplots were created in R and linear regression models were fitted to each plot. Mantel tests (Sokal and Rohlf 1995; Wagner and Fortin 2015) were used to assess the correlations between genetic distance and either geographic or environmental distances. The significance of these relationships was determined using 9999 permutations.

A mixed model was utilized to analyze the relationships between genetic distance, geographic distance, and environmental distance. To address potential collinearity issues that could create confounding results in the mixed model, variance inflation factor (VIF) assessments (O’brien 2007) were conducted to confirm low multicollinearity for each of the three distance matrices (Supplemental Table 14).

To complement this analysis, a two-dimensional kernel density calculation was applied to visualize the concentration of data points within the scatterplots. To discern potential non-linear relationships between the distance matrices, a locally estimated scatterplot smoothing technique, implemented via the loess.smooth() function in R (Gareth et al. 2013), was employed. This nonlinear fit was compared against the linear model, with R² scores for both models calculated to assess and contrast their respective fits.

### Climate Data Extraction

Climate data was extracted from the ERA5-Land monthly averaged dataset provided by the Copernicus Climate Change Service (Copernicus Climate Change Service 2019). The original dataset was characterized by a temporal resolution of 1 hour and a native spatial resolution of 9 km on a reduced Gaussian grid (TCo1279). To facilitate broader accessibility and suitability for diverse analyses, the data underwent regridding to a regular lat-lon grid with a finer resolution of 0.1x0.1 degrees.

In this study, we initially extracted a total of 13 environmental variables for comprehensive analysis. These variables encompass a diverse range of climate parameters: 10m eastward wind, 10m northward wind, 2m temperature, evaporation from bare soil, leaf area index for high vegetation, leaf area index for low vegetation, water retention capacity of land, snowfall, surface net solar radiation, surface runoff, total evaporation, total precipitation, and volumetric soil water layer 1. Table 3 provides detailed descriptions of each variable.

**Table 3:**
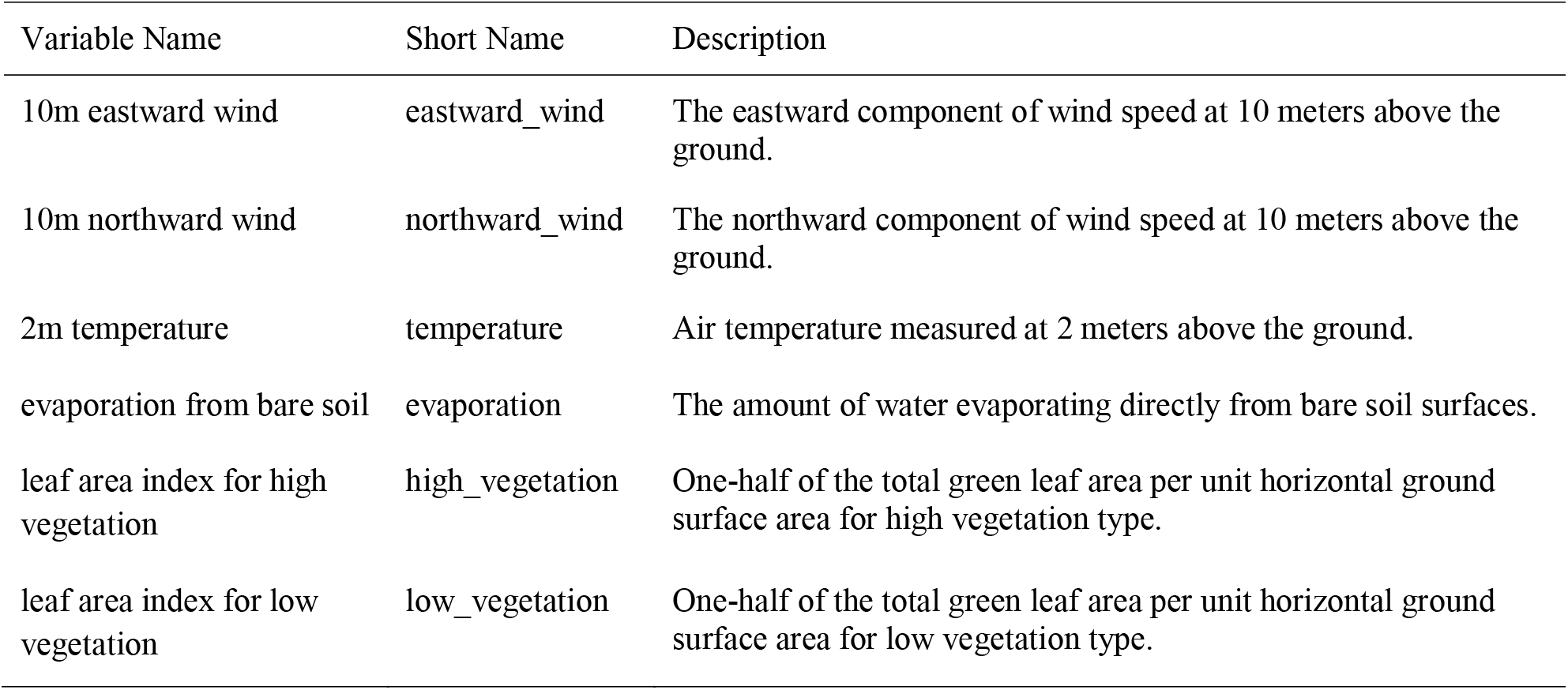
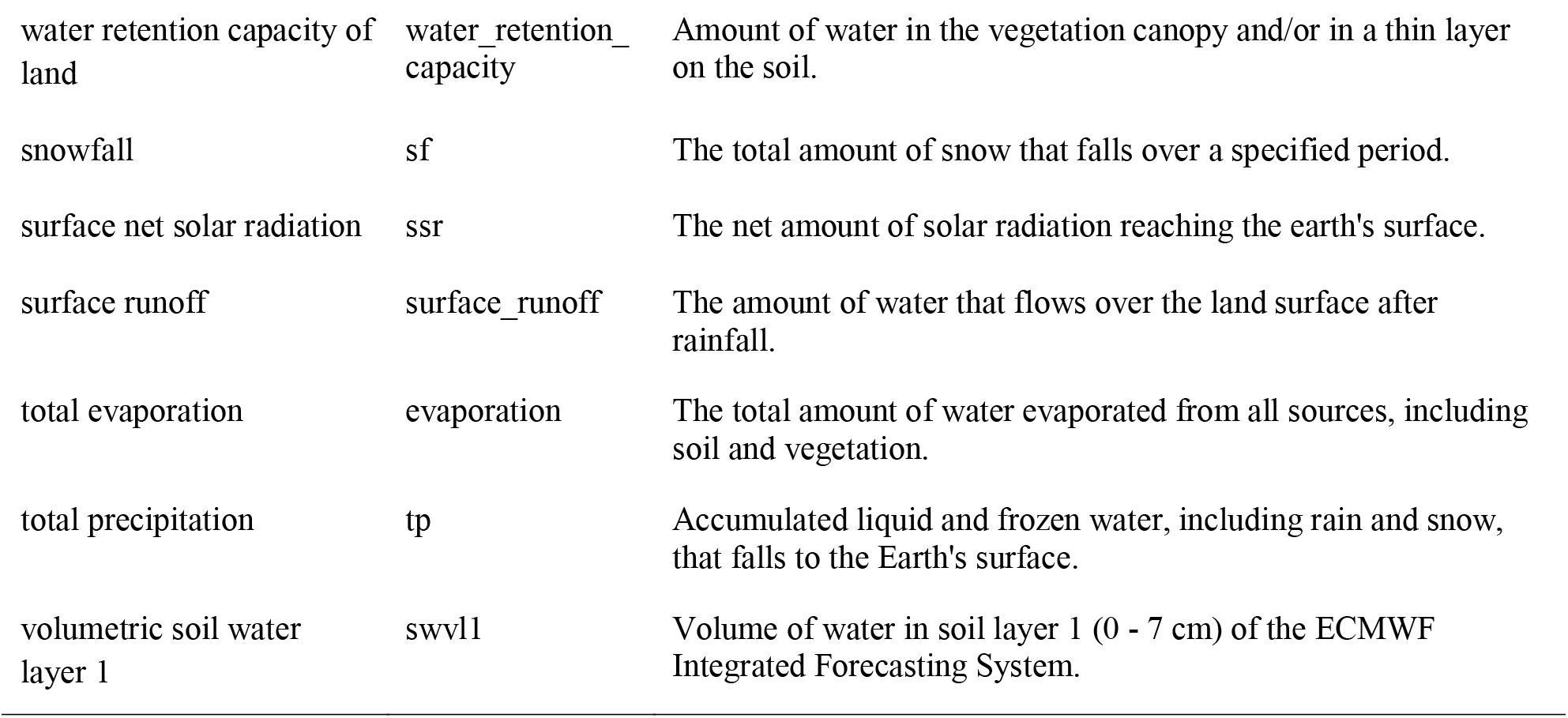
Description of environmental variables.

| Variable Name | Short Name | Description |
| --- | --- | --- |
| 10m eastward wind | eastward_wind | The eastward component of wind speed at 10 meters above the ground. |
| 10m northward wind | northward_wind | The northward component of wind speed at 10 meters above the ground. |
| 2m temperature | temperature | Air temperature measured at 2 meters above the ground. |
| evaporation from bare soil | evaporation | The amount of water evaporating directly from bare soil surfaces. |
| leaf area index for high vegetation | high_vegetation | One-half of the total green leaf area per unit horizontal ground surface area for high vegetation type. |
| leaf area index for low vegetation | low_vegetation | One-half of the total green leaf area per unit horizontal ground surface area for low vegetation type. |
| water retention capacity of land | water_retention_capacity | Amount of water in the vegetation canopy and/or in a thin layer on the soil. |
| snowfall | sf | The total amount of snow that falls over a specified period. |
| surface net solar radiation | ssr | The net amount of solar radiation reaching the earth's surface. |
| surface runoff | surface_runoff | The amount of water that flows over the land surface after rainfall. |
| total evaporation | evaporation | The total amount of water evaporated from all sources, including soil and vegetation. |
| total precipitation | tp | Accumulated liquid and frozen water, including rain and snow, that falls to the Earth's surface. |
| volumetric soil water layer 1 | swvl1 | Volume of water in soil layer 1 (0 - 7 cm) of the ECMWF Integrated Forecasting System. |

Prior to processing this complex dataset, steps were taken to minimize variable redundancy. This involved evaluating the variance for each variable across all samples, examining pairwise correlation coefficients, and assessing the distribution of each variable. Variables with a total variance across samples of zero, a pairwise correlation coefficient exceeding 0.70, or showing extreme distribution patterns were identified and eliminated. This refinement process led to the removal of 5 variables: surface net solar radiation, total evaporation, total precipitation, volumetric soil water layer 1, and snowfall. Consequently, the variable set was reduced to 8 environmental variables for all subsequent analyses (Supplemental Figure 7).

The reduced dataset was processed using Python scripts utilizing the xarray and pandas libraries. These scripts calculated annual averages for the 8 remaining climate variables for the year 2012. The data processing was specifically tailored to extract meaningful climate insights from NetCDF format files from the ERA5-Land monthly averaged dataset, with each variable’s average computed based on its geographical location coordinates. This approach aimed to transform the extensive climate data into a format suitable for later Genome-Environment Association (GEA) (Kamvar et al. 2017) analyses.

### Identification of Genotype-Environment Associations

To explore potential adaptive divergence in *Cx. tarsalis* populations, we employed two different approaches: a genome-environment association (GEA) (Kamvar et al. 2017) via latent factor mixed models (LFMM) (Frichot et al. 2013; Caye et al. 2019)(Sokal and Rohlf 1995; Wagner and Fortin 2015) and a redundancy analysis (RDA) (van den Wollenberg 1977) implemented in the vegan package in R (Oksanen et al. 2019).

The LFMM approach utilizes a univariate testing framework, modeling each SNP and environmental variable using the lfmm_test() function from the LFMM package in R (Caye et al. 2019). We initially conducted a Principal Component Analysis (PCA) on the environmental variables, focusing on the first principal component—a linear combination of the 8 climate variables—as the predictor in the LFMM. We also consider the inclusion of the second and third principal components as additional predictors in a subsequent LFMM, as detailed in the Supplemental Materials (Supplemental Figure 8 and Supplemental Table 15 - 17)

In assessing the association between SNPs and environmental variables using the LFMM, we evaluated the Genomic Inflation Factor (GIF) to determine the model’s efficacy in handling potential confounding factors. In our analysis, a Genomic Inflation Factor (GIF) value of 1.23 was used to adjust the p-values for potential inflation due to confounding variables, a process that is automatically incorporated within the output of the lfmm_test() function in R (Caye et al. 2019). Subsequently, we converted these GIF-adjusted p-values to q-values using the qvalue() function in R. This conversion is to refine the significance thresholds for individual tests, particularly under the framework of multiple hypothesis testing. It enhances the precision of FDR control, crucial in large-scale testing scenarios like ours. We then employed FDR control measures using these q-values, identifying candidate results falling below our predefined FDR threshold of 0.1.

The RDA operates as a multifaceted ordination technique, evaluating multiple loci concurrently with environmental variables. For this analysis, the significance of both the overall RDA model and its individual constrained axes was assessed. This assessment utilized the anova.cca() function from the vegan package in R (Oksanen et al. 2019; Borcard et al.), facilitating a comprehensive examination of the null hypothesis (an absence of a linear relationship between SNP data and environmental variables). To select candidate SNPs for local adaptation, we identified SNPs that significantly deviated from the mean loadings, using a threshold of 3 standard deviations. Statistical significance was determined based on p-values below the threshold of 0.001.

To visualize results and identify relationships between RDA components and other factors, we utilized the vegan package in R (Oksanen et al. 2019; Borcard et al.)to generate RDA tri-plots between each pair of the RDA components. To enhance the interpretability of our ordination plots, we employed symmetrical scaling (Borcard et al.). This scaling method adjusts the SNP and individual scores by the square root of the eigenvalues, providing a clearer representation of the relationships between variables and samples.

### Detecting SNPs Under Selection

To find which SNPs that were significantly associated with environmental variables also showed signatures of natural selection, we performed a Bayesian selection inference as implemented in BayeScan (Foll and Gaggiotti 2008; Foll et al. 2010; Fischer et al. 2011). We also independently identified outliers using a PCA-based method implemented in the pcadapt R package (Privé et al. 2020), and then filtered for common candidate SNPs that were identified as significant across 4 different analyses: LFMM, RDA, BayeScan, and PCAdapt.

## Data Availability

All custom scripts used in this study are available at GitHub Repository: https://github.com/Afei99357/Culex_Tarsalis_GWAS_manuscript.git.

The data used in the GitHub code is available at: Dryad Data Repository: https://doi.org/10.5061/dryad.51c59zwh3.

The *Cx. tarsalis* whole genome sequencing data and the raw sample RAD-Seq data are stored in the Sequence Read Archive (SRA). The accession number for the whole genome is in project PRJNA1124366, and the RAD-Seq data is in project PRJNA1126219 (https://dataview.ncbi.nlm.nih.gov/object/PRJNA1126219).

## Acknowledgements

The authors would like to thank E. Prates, M. Shah, M. Pavicic and M. Cashmann for their insights on protein-coding changes in the *Cx. tarsalis* genome. The authors would also like to thank P. Lado for her insights on an earlier version of this manuscript. Finally, the authors wish to acknowledge the University Research Computing group at UNC Charlotte for providing access to the HPC resources needed for this research project.

This research was supported by a cooperative agreement with the United States Department of Agriculture (NACA 58-3022-1-002).

