## Supplemental Figure for "Climate Adaptation and Genetic Differentiation in the Mosquito Species *Culex tarsalis*"

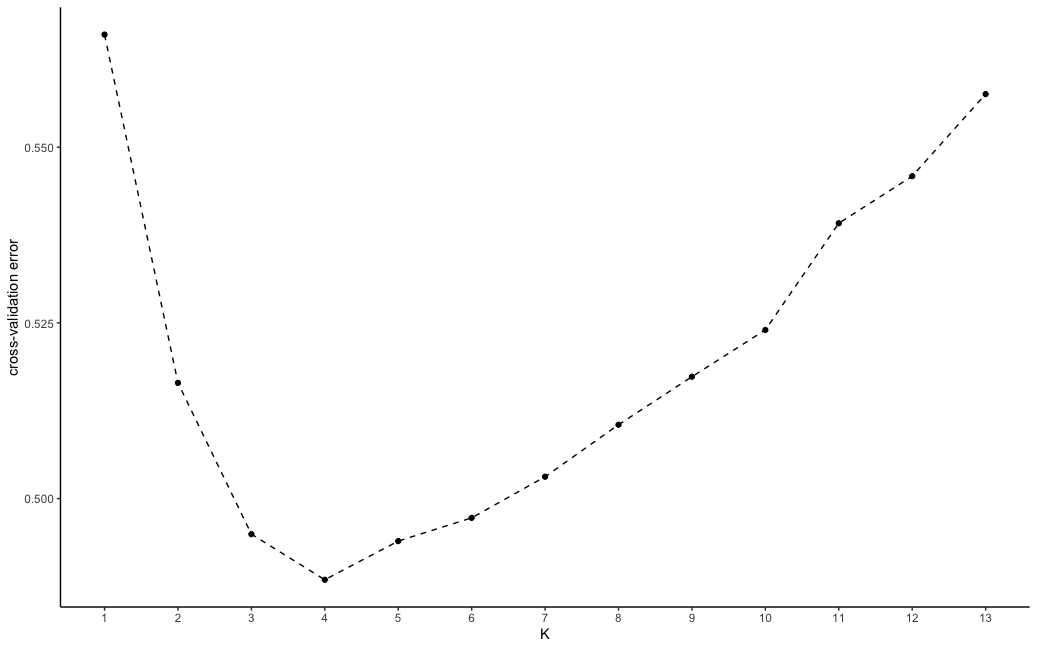

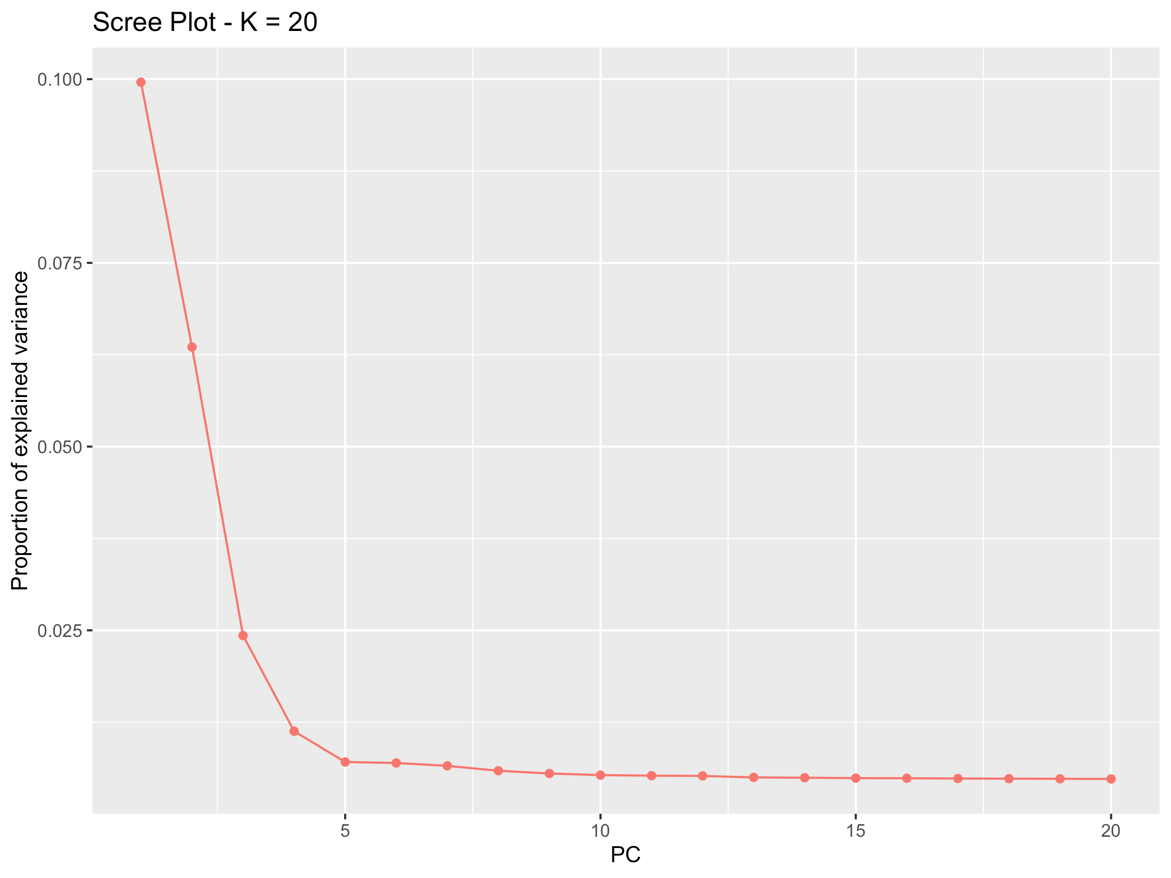

**Supplemental Figure 1: Optimal number of Clusters (K).** Top plot shows the cross-validation error estimated for K values ranging from 1 to 13. Bottom plot indicates the proportion of variance explained in a PCA of SNP data.

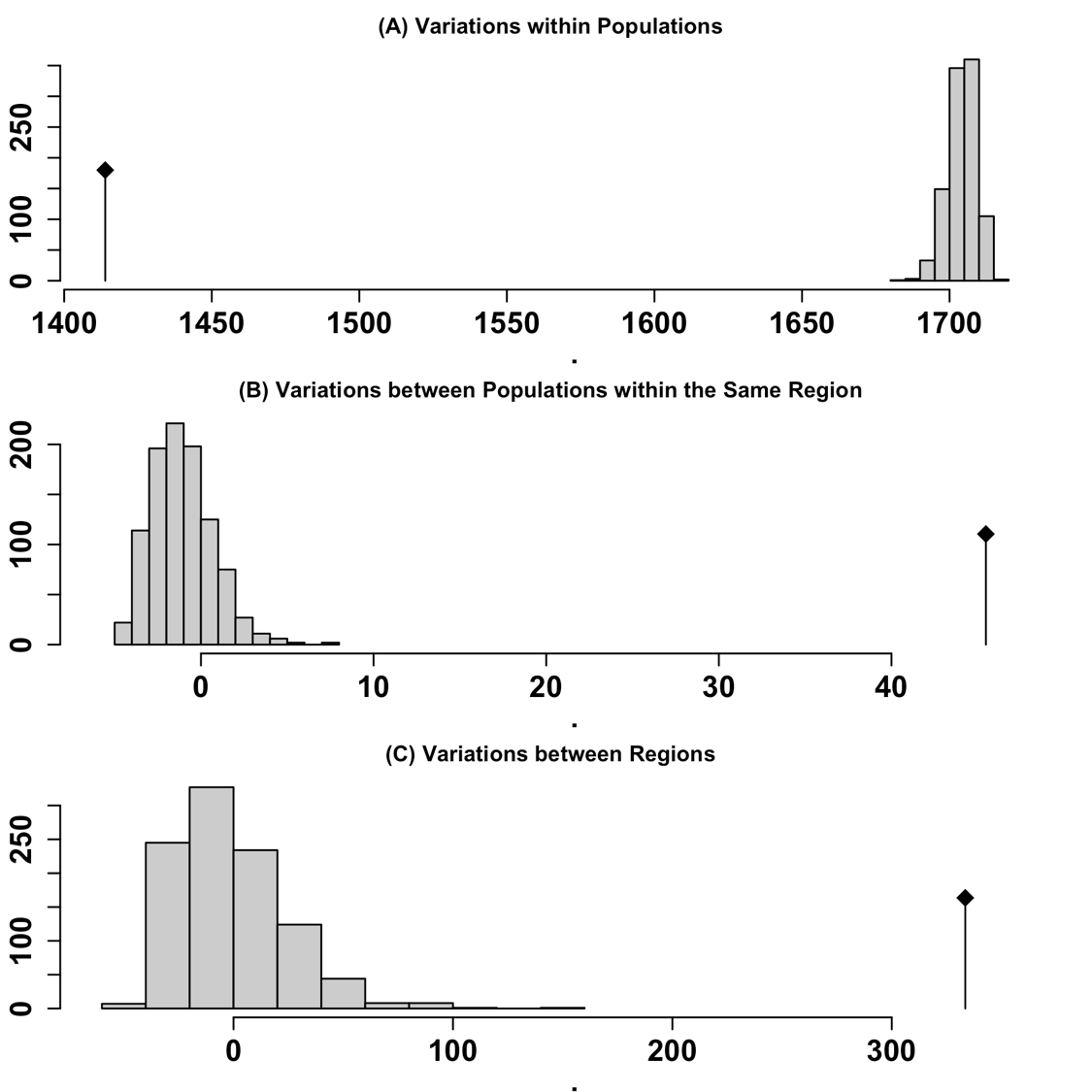

**Supplemental Figure 2: Analysis of Molecular Variance (AMOVA).** Gray bars indicate the expected distributions based on 999 random permutations, while black bars indicate the observed phi values. (A) Variation within populations, (B) Variation between populations within the same region, and (C) Variation between different regions.

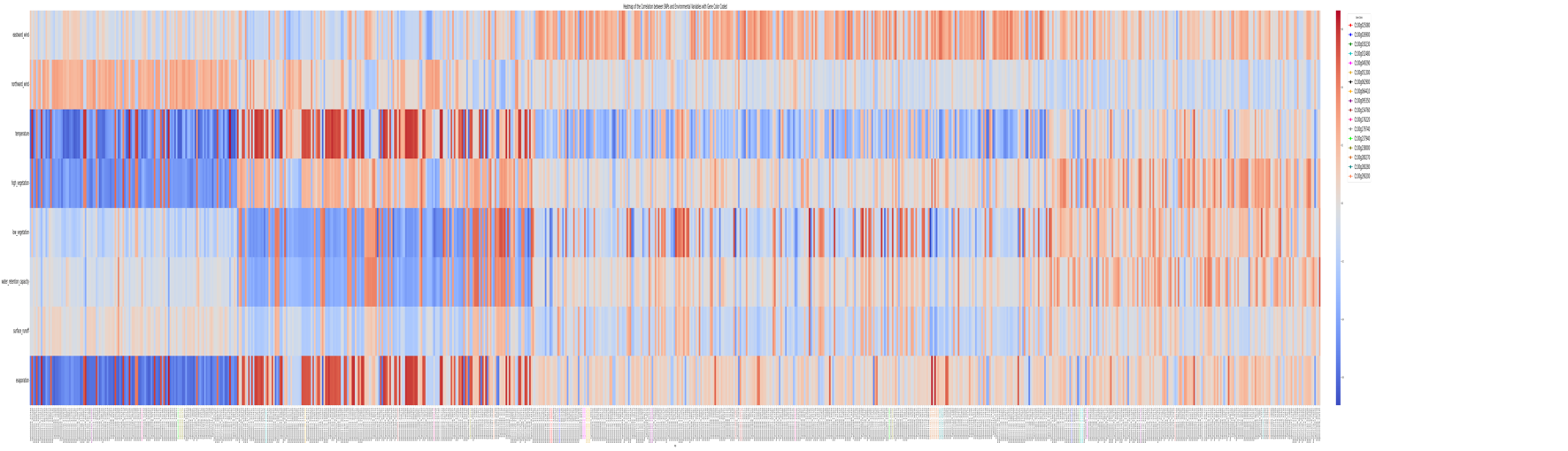

**Supplemental Figure 3: Correlation Heatmap between Environmental Variables and RDA Candidate SNPs.** This heatmap illustrates the correlation between the environmental variables and candidate SNPs identified through the first four constrained axes of Redundancy Analysis (RDA). Each column on the x-axis represents a candidate SNP. Detailed SNP names and associated information on the 17 genes identified by SIFT4G

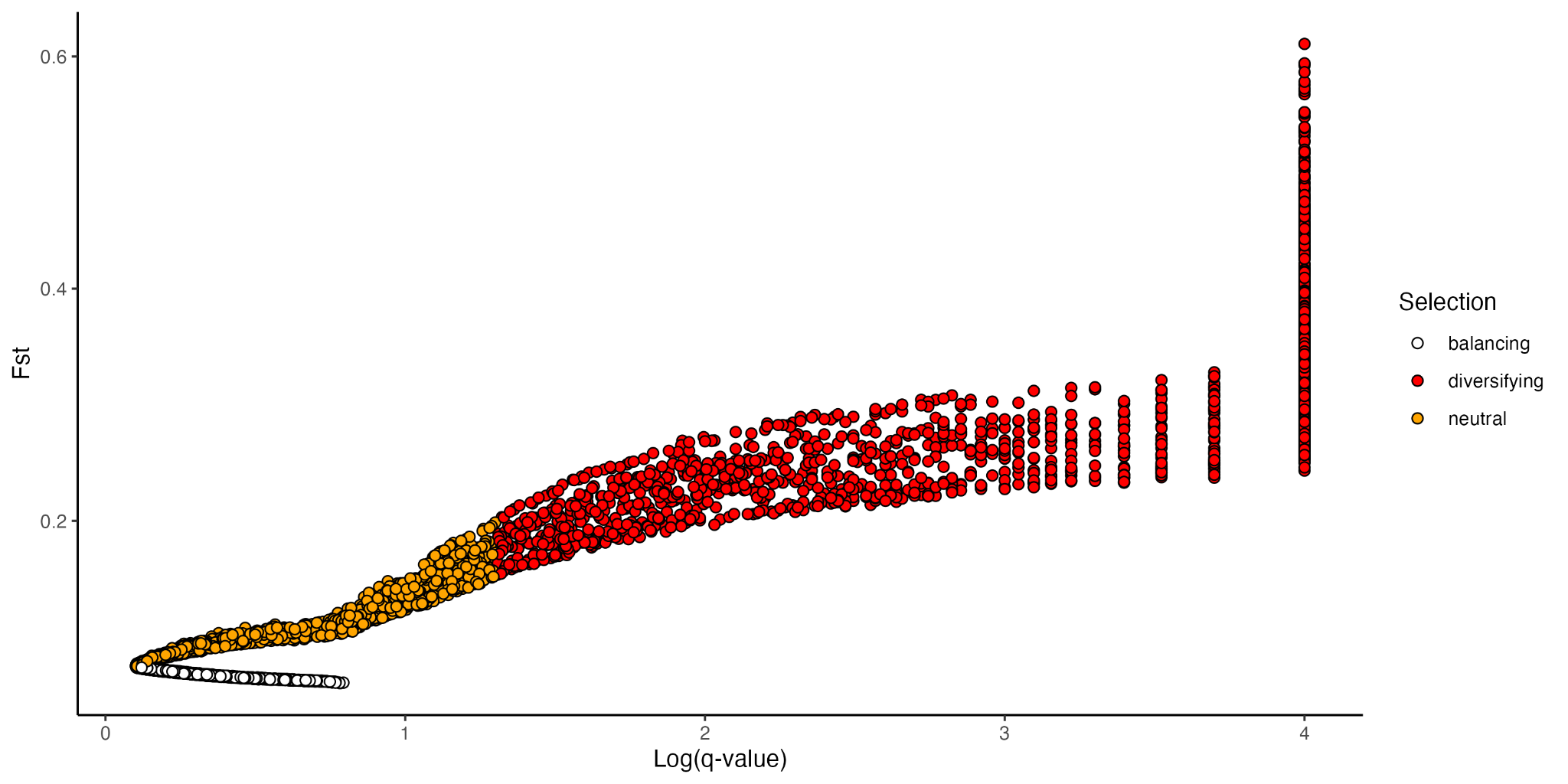

**Supplemental Figure 4: BayeScan results.**

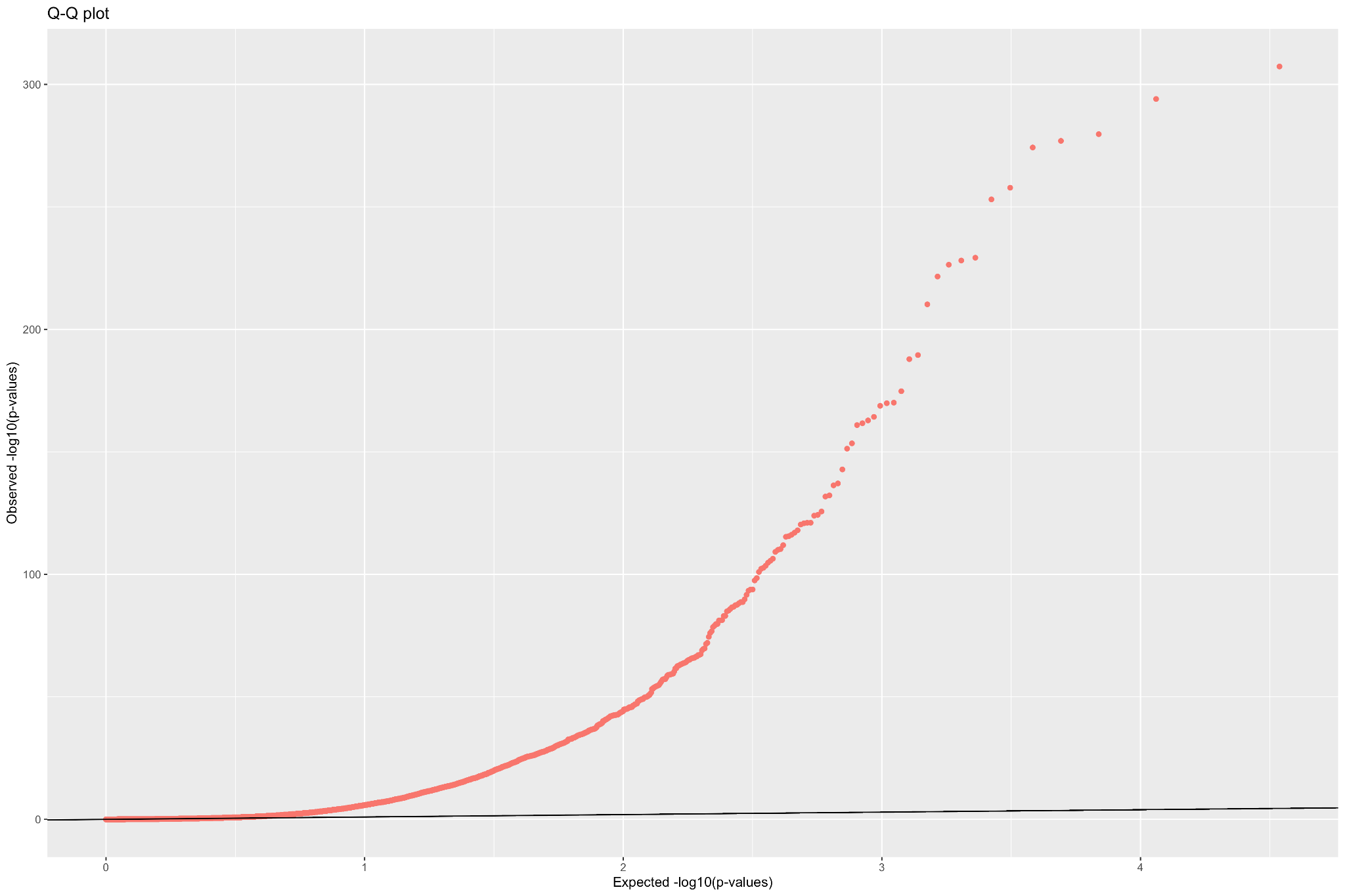

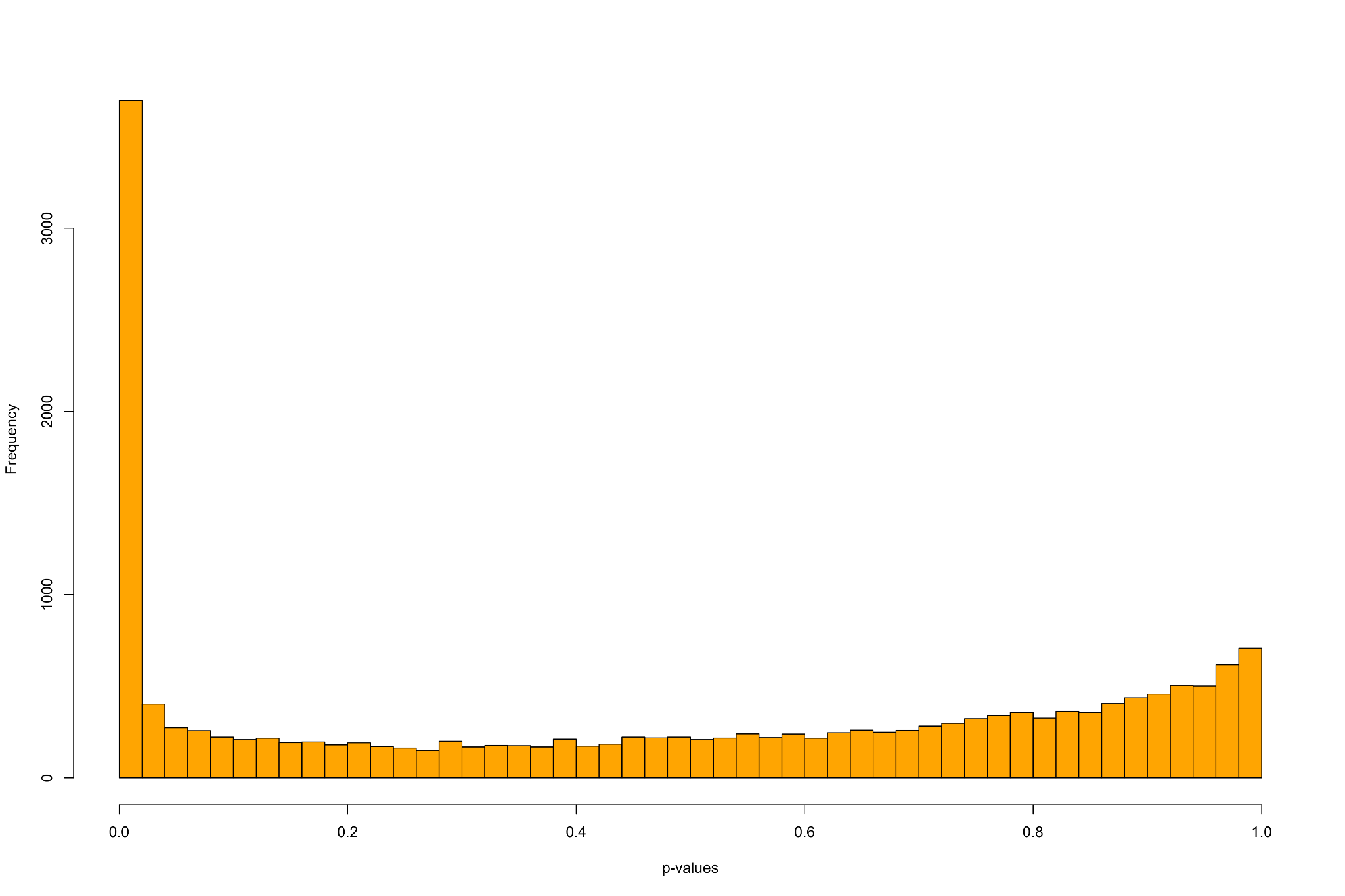

**Supplemental Figure 5: PCAdapt results.**

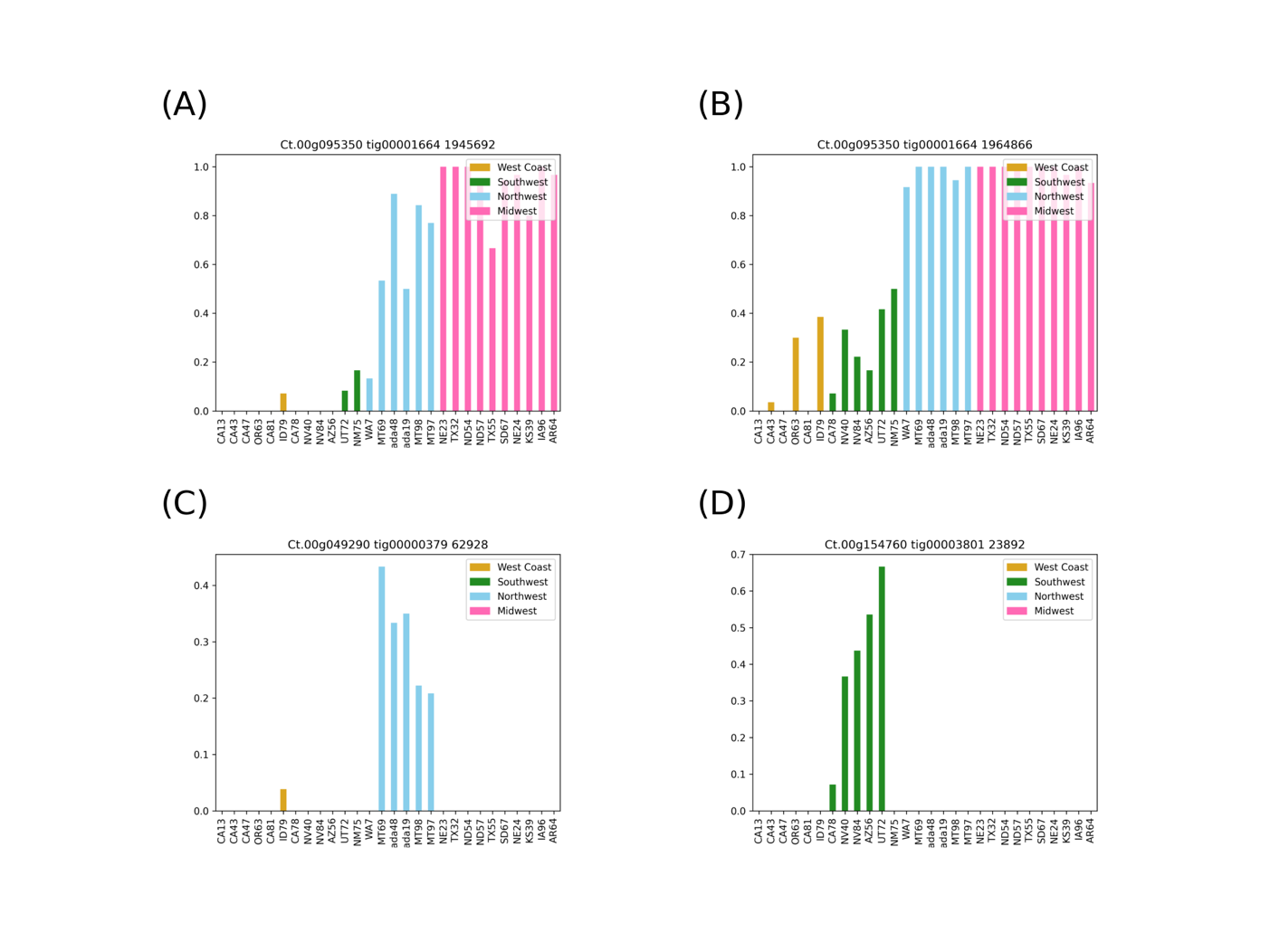

**Supplemental Figure 6. Allele frequency distribution of selected candidate SNPs**

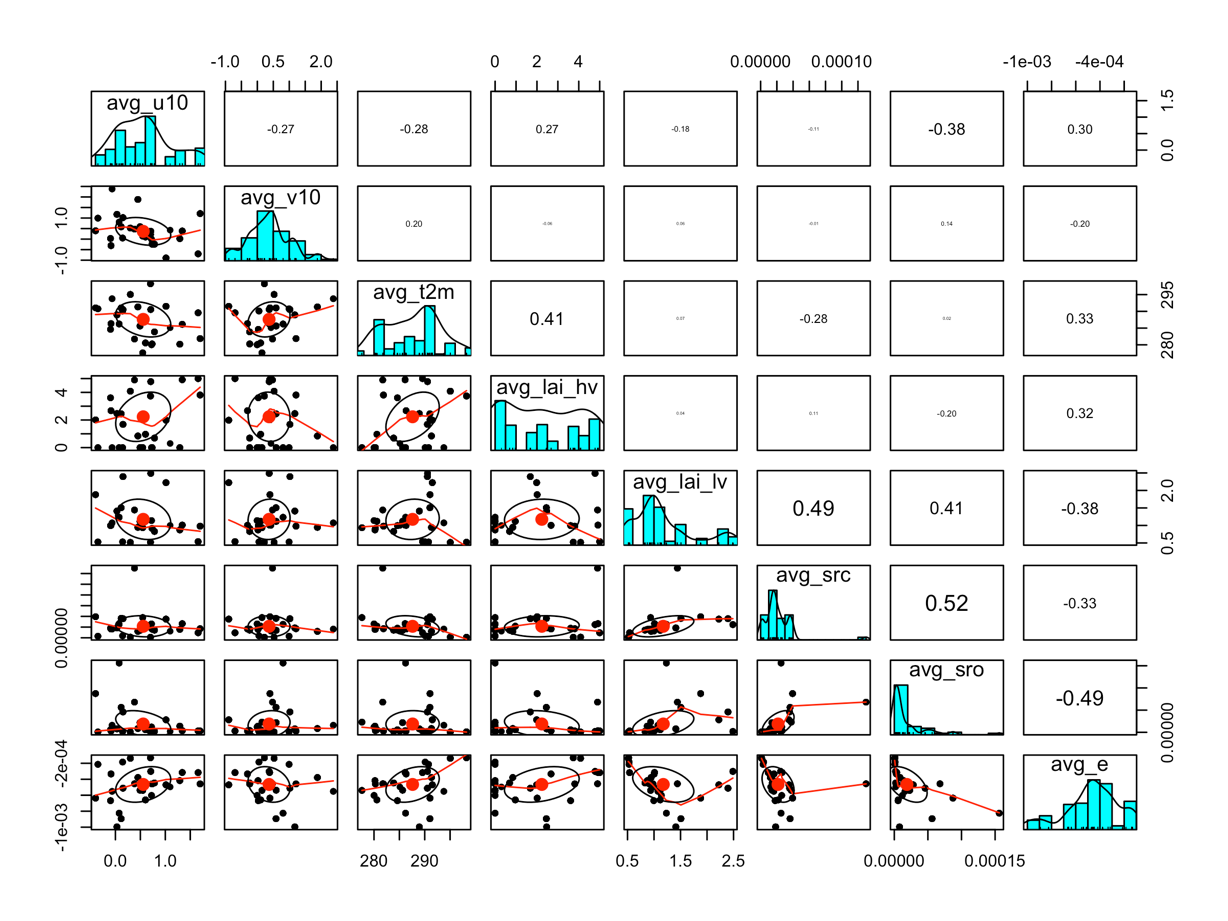

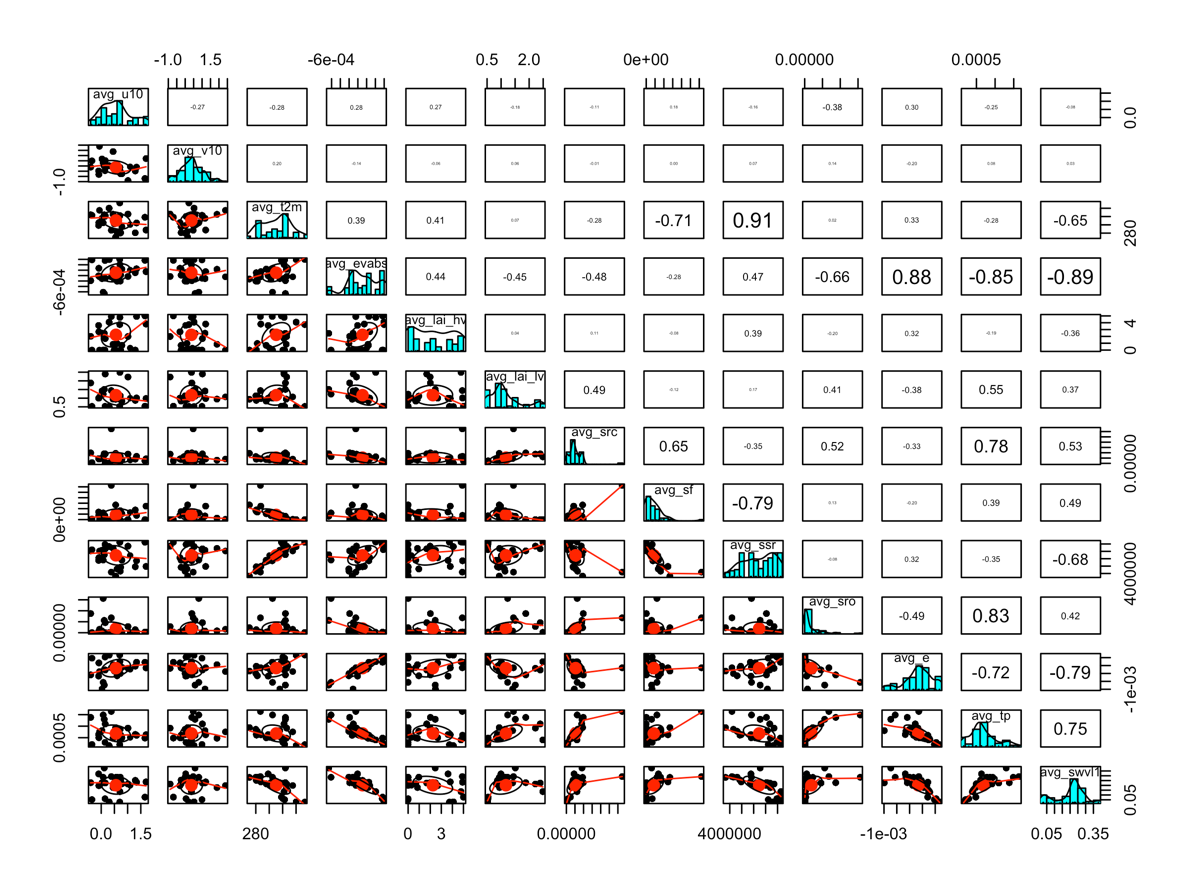

**Supplemental Figure 7: Climate variable correlations and distributions.** Top panel shows the pairwise correlations and distributions for all 13 environmental variables downloaded from the Copernicus Climate Change Service. Bottom panel shows the correlations and distributions for the 8 remaining variables after removing variables with low variance or correlations >0.70.

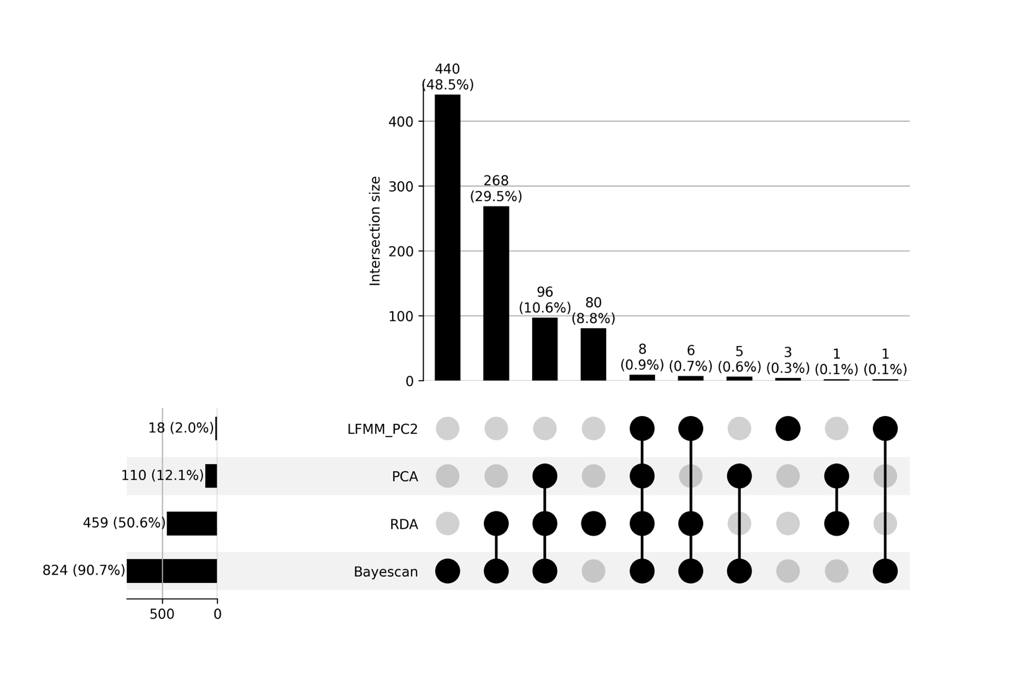

(A)

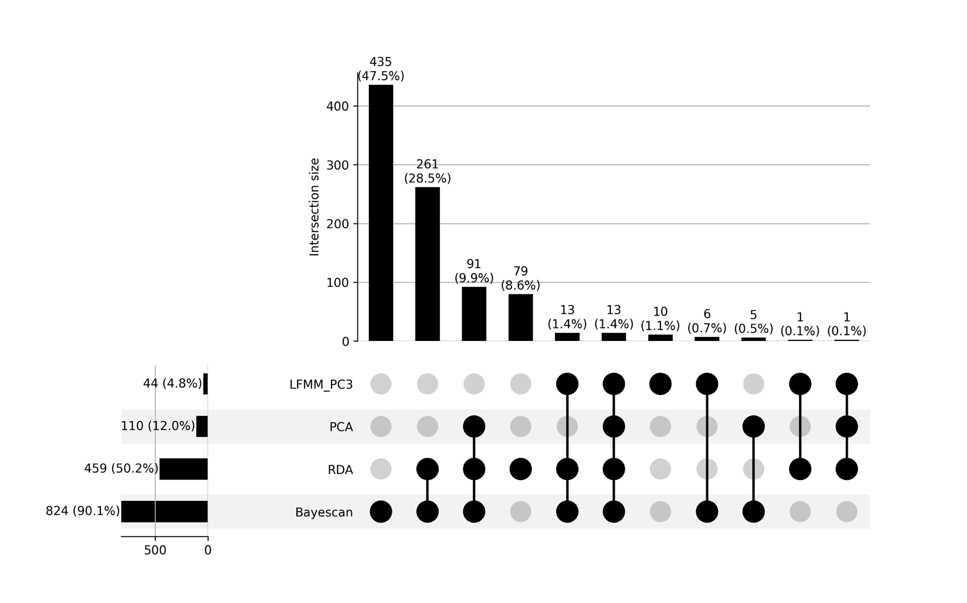

(B)

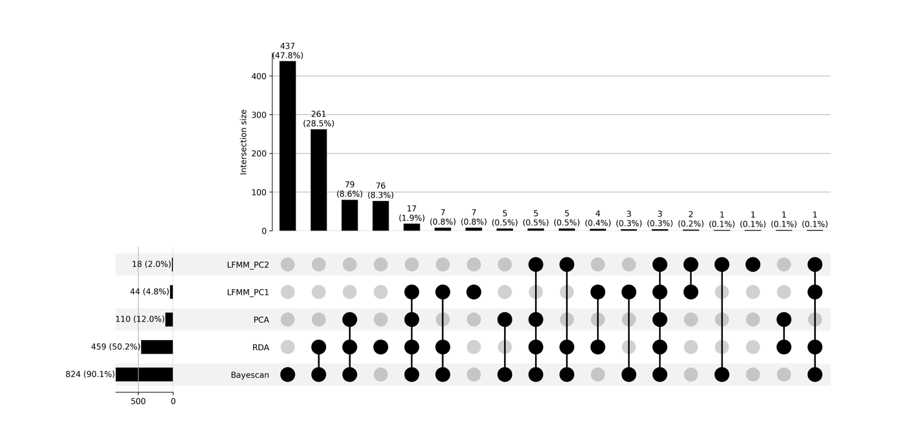

(C)

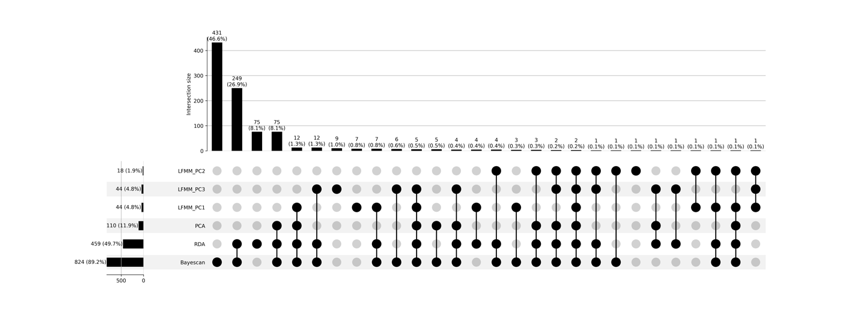

(D)

**Supplemental Figure 8:** Upset Plot of Candidate SNPs Identified for Local Adaptation and Environmental Association Across LFMM, RDA, Pcadapt, and BayesScan Analysis in Cx. Tarsalis. (A) LFMM PC2 across other methods (B) LFMM PC2 across other methods

(C) LFMM PC1 and PC2 across other methods (D) LFMM PC1, PC2 and PC3 across other methods
